# Perceiving structure in unstructured stimuli: Implicitly acquired prior knowledge impacts the processing of unpredictable transitional probabilities

**DOI:** 10.1101/738419

**Authors:** Andrea Kóbor, Kata Horváth, Zsófia Kardos, Dezso Nemeth, Karolina Janacsek

## Abstract

It is unclear how implicit prior knowledge is involved and remains persistent in the extraction of the statistical structure underlying sensory input. Therefore, this study investigated whether the implicit knowledge of 2^nd^ order transitional probabilities characterizing a stream of visual stimuli impacts the processing of unpredictable transitional probabilities embedded in a similar input stream. Young adults (*N* = 50) performed a four-choice reaction time (RT) task that consisted of structured and unstructured blocks. In the structured blocks, more probable and less probable short-range nonadjacent transitional probabilities were present. In the unstructured blocks, the unique combinations of the short-range transitional probabilities occurred with equal probability; therefore, they were unpredictable. All task blocks were visually identical at the surface level. While one-half of the participants completed the structured blocks first followed by the unstructured blocks, this was reversed in the other half of them. The change in the structure was not explicitly denoted, and no feedback was provided on the correctness of each response. Participants completing the structured blocks first showed faster RTs to more probable than to less probable short-range transitional probabilities in both the structured and unstructured blocks, indicating the persistent effect of prior knowledge. However, after extended exposure to the unstructured blocks, they updated this prior knowledge. Participants completing the unstructured blocks first showed the RT difference only in the structured blocks, which was not constrained by the preceding exposure to unpredictable stimuli. The results altogether suggest that implicitly acquired prior knowledge of predictable stimuli influences the processing of subsequent unpredictable stimuli. Updating this prior knowledge seems to require a longer stretch of time than its initial acquisition.

## 1. Introduction

Acquiring implicit knowledge of the statistical structure organizing environmental events is crucial for many cognitive functions and contributes to the automatization of behaviors (Armstrong, Frost, & Christiansen, 2017; Aslin, 2017; Kaufman et al., 2010; Maheu, Dehaene, & Meyniel, 2019). This ability involves not only the mere extraction of various statistical structures but also the efficient use of the acquired implicit knowledge across situations that differ in specific features at the surface level but share common features at the structural level. In everyday life, if conditions are substantially similar, we usually learn fast how to use the updated versions of applications or operating systems without checking manuals, running online searches, or even consciously accessing the course of our actions by using previous experiences. However, the potential stability of the already acquired implicit knowledge when applied in similar situations has not been completely elucidated (Bulgarelli & Weiss, 2016; Conway, 2020; R. Frost, Armstrong, & Christiansen, 2019). Therefore, we investigate the stability of implicit knowledge of a statistical structure underlying a stream of visual stimuli that remains the same at the surface level but, in time, becomes unpredictable at the structural level.

According to the broad frameworks of cognitive processing, learning, and decision making, the processing of new information and the formation of expectations about future events are guided by inferences based on prior experiences (e.g., Daw, Gershman, Seymour, Dayan, & Dolan, 2011; Friston, 2005; Friston, 2010; Friston, Stephan, Montague, & Dolan, 2014; Griffiths, Kemp, & Tenenbaum, 2008; Shohamy & Daw, 2015). This also pertains to randomness perception (Hahn & Warren, 2009; Sun et al., 2015; Sun & Wang, 2010; Teigen & Keren, 2020; Warren, Gostoli, Farmer, El-Deredy, & Hahn, 2018), binary choice behavior (Feher da Silva & Baldo, 2012; Gaissmaier & Schooler, 2008; James & Koehler, 2011) as well as implicit statistical learning (Conway, 2020; Qian, Jaeger, & Aslin, 2012). The persistence of the primarily learned statistical structure and its influence on further processing have been evidenced by behavioral (e.g., Bulgarelli & Weiss, 2016; Gebhart, Aslin, & Newport, 2009; Lany, Gómez, & Gerken, 2007) as well as neurocognitive measures (e.g., Honbolygó & Csépe, 2013; Karuza et al., 2016; Mullens et al., 2014; Todd, Frost, Fitzgerald, & Winkler, 2020; Todd, Provost, & Cooper, 2011). However, statistical structures can differ in characteristics and complexity (Conway, 2020), and multiple statistical structures can be acquired even from the same stimulus sequence (Conway & Christiansen, 2001; Daltrozzo & Conway, 2014; R. Frost et al., 2019; Kóbor et al., 2018; Simor et al., 2019).

According to the model proposed by Meyniel, Maheu, and Dehaene (2016), instead of simpler statistics such as frequencies and alternations of events, the computation of time-varying, non-stationary, local transitional probabilities between consecutive events could be considered as the “building block” of implicit statistical learning and knowledge (see also Maheu et al., 2019; Orbán, Fiser, Aslin, & Lengyel, 2008). Humans have been found to be highly proficient in extracting even the nonadjacent transitional probabilities, referring to predictive relations between elements of a sequence that includes ordered stimuli interspersed with random ones (Conway, 2020; R. L. A. Frost & Monaghan, 2016; Malassis, Rey, & Fagot, 2018; Mueller, Milne, & Männel, 2018; Rey, Minier, Malassis, Bogaerts, & Fagot, 2018).

Using transitional probabilities in a series of experiments, Gebhart et al. (2009) changed the underlying statistical structure of stimuli in the middle of an auditory statistical learning task. They successively presented two different but overlapping artificial speech streams composed of trisyllabic nonsense words characterized by transitional probabilities. In this way, the surface of the stimuli remained similar throughout the task while their structure changed. If the change was not explicitly signaled or the second structure was not presented for a tripled duration, participants only learned the first structure. This indicated that the primarily experienced structure limited the capacity to acquire the successive structure. However, in this study, a certain statistical structure determined by transitional probabilities was always present during the task (see also Bulgarelli & Weiss, 2016; Weiss, Gerfen, & Mitchel, 2009; Zinszer & Weiss, 2013). Therefore, it is unclear whether the results would have been the same if the statistical structure per se had been eliminated. For instance, it could be clarified whether changing only the predictability of the same nonadjacent transitional probabilities over the course of learning influences their later extraction.

Furthermore, in the study by Gebhart et al. (2009), after exposure to the speech stream, knowledge of the statistical structures was measured with two-alternative forced-choice test trials in which familiarity judgments were provided. Meanwhile, processing-based or “online” tasks should be favored, since these tasks more likely reflect implicitly acquired statistical knowledge about which no consciously accessible representations are available. These tasks could also capture the trajectory of acquisition and provide information about the stability of the underlying processes when these processes actually operate (Christiansen, 2018; R. Frost et al., 2019). Therefore, it remains to be tested with an online, unsupervised statistical learning task (Fiser & Aslin, 2001; Qian et al., 2012) how changing the predictability of nonadjacent transitional probabilities impacts further acquisition.

Consequently, in the present study, we used a four-choice reaction time (RT) task to online measure the implicit processing and acquisition of a sequence composed of 2^nd^ order nonadjacent transitional probabilities. In this sequence, elements in position *n* – 2 predicted elements in position *n* with high or low probability. Unknown to participants, half of the task blocks included an alternating regularity between nonadjacent trials, yielding more probable and less probable short-range transitional probabilities (see Fig. 1). The short-range transitional probabilities were three successive trials, hereafter referred to as triplets. In the other half of the task blocks, the alternating regularity was absent, and unique triplets appeared with equal probability. The task blocks were labeled as structured and unstructured blocks, according to the presence or absence of the alternating regularity. By creating either biased (high or low) or equal probabilities of triplets, stimuli were predictable in the structured blocks and unpredictable in the unstructured blocks. With this design, it could be tested how prior knowledge of the predictability of triplets influences their processing when predictability changes from the first to the second half of the task. To this end, while one-half of the fifty participants completed the structured blocks first followed by the unstructured blocks, the other half of the participants completed the unstructured blocks first followed by the structured blocks. Participants of both groups received neither explicit information on the midstream change in structure nor feedback on the correctness of each response throughout the task.

**Figure 1.**
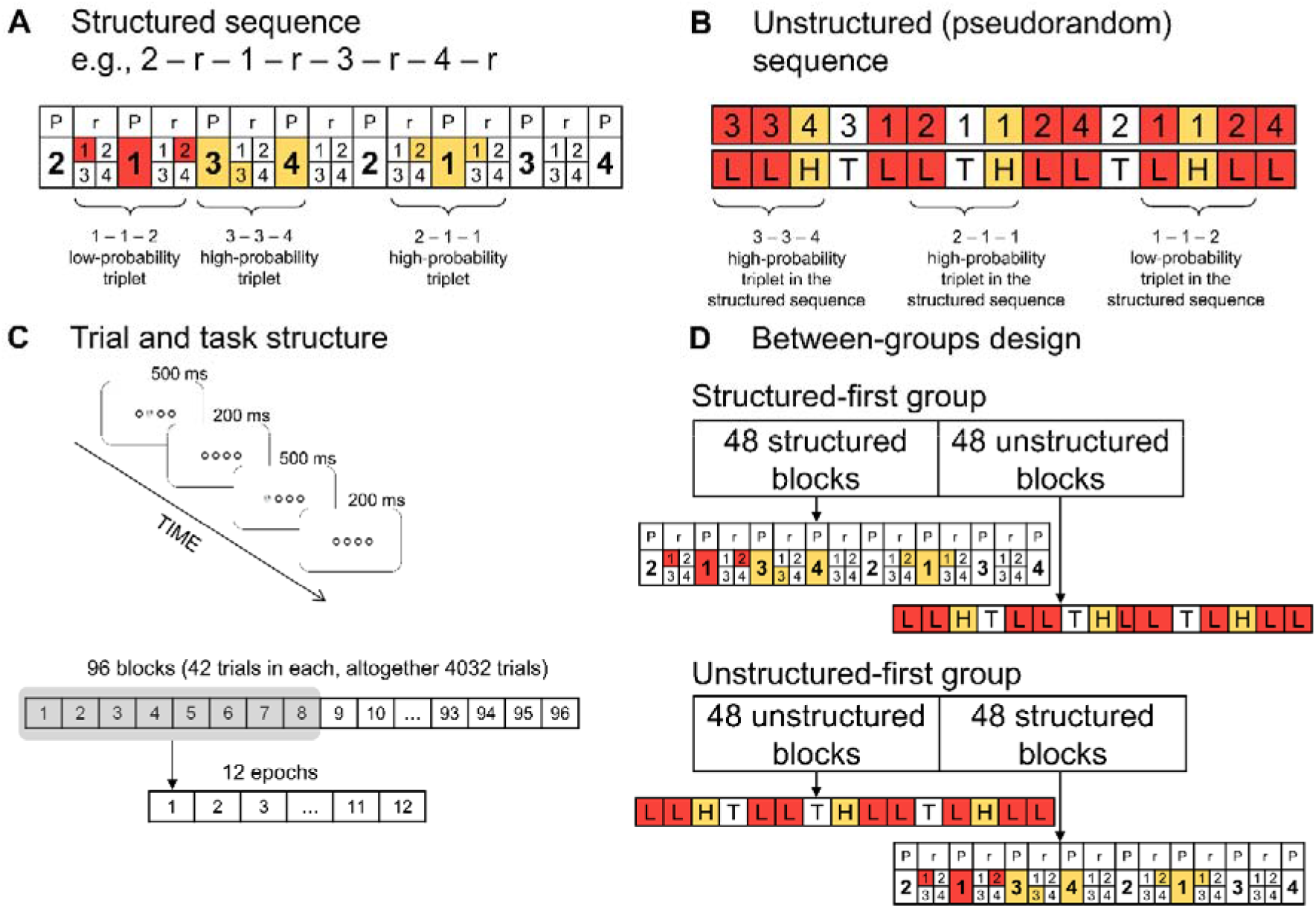
Design of the experiment. (A) The presentation of stimuli in the structured ASRT sequence followed an eight-element regularity, within which pattern (P) and random (r) elements alternated with one another. Numbers denote the four different stimulus positions on the screen. The alternating regularity made some runs of three consecutive trials (triplets) more probable than others. High-probability triplets are denoted with gold shading and low-probability triplets are denoted with coral shading. (B) From the unstructured (pseudorandom) sequence, the alternating regularity was omitted, but the same unique triplets as in the structured sequence appeared with equal probability. Note that the probability of triplets only differs in the structured sequence and their probability is equal in the unstructured sequence. However, triplets in the unstructured sequence are still labeled as either high- or low-probability according to their actual probability in the structured sequence. Gold shading (upper row) and capital letter “H” (lower row) denote the third element of high-probability triplets, coral shading and “L” denote the third element of low-probability triplets, while white shading and “T” as “trill” denote the third element of some of those low-probability triplets that were eliminated from the analyses (see Statistical analysis section). Numbers denote the four different stimulus positions on the screen. Note that each stimulus (trial) is categorized as either the third element of a high- or a low-probability triplet in both sequences. For a given participant, at the level of unique triplets, the high- and low-probability triplets are the same in the structured and unstructured sequences. (C) In this version of the task, a stimulus appeared in one of four horizontally arranged empty circles on the screen in every 700 ms. Participants had to respond with one of the four response keys that corresponded to the position of the stimulus. They completed altogether 96 blocks, and eight-block-long units of the task were collapsed into larger time bins labeled as epochs. (D) While the Structured-first group (*n* = 25) completed 48 structured blocks followed by 48 unstructured blocks, the Unstructured-first group (*n* = 25) completed 48 unstructured blocks followed by 48 structured blocks.

If the influence of the biased probabilities acquired over the structured blocks persisted throughout the task, the RT difference between the more probable and less probable triplets would be similar across the structured and unstructured blocks for participants completing the structured blocks first. Moreover, it could also be explored how long the influence of this already acquired knowledge would last. Meanwhile, prior knowledge of equal probabilities emerging over the unstructured blocks could also persist and influence the further acquisition of biased probabilities. Accordingly, for participants completing the unstructured blocks first, no RT difference between the triplets is expected over the unstructured blocks. The RT difference over the structured blocks would emerge only in a slower, more gradual manner (cf. Zhao et al., 2019). However, as the lack of RT difference could persist throughout the task, it is also conceivable that these participants would not acquire the biased probabilities over the structured blocks.

## 2. Material and methods

### 2.1 Participants

Fifty healthy young adults took part in the experiment.^1^ They were undergraduate students from Budapest, Hungary. Participants had normal or corrected-to-normal vision, and according to the predefined inclusion criteria, none of them reported a history of any neurological and/or psychiatric condition, and none of them was taking any psychoactive medication. Half of the participants were randomly assigned to the Structured-first group (*n* = 25), while the other half was assigned to the Unstructured-first group (*n* = 25). The groups were differentiated by which half of the experimental task they started with; this is explained in the Procedure section below. Descriptive characteristics of participants in the two groups and their performance on standard neuropsychological tests are presented in Table 1. All participants provided written informed consent before enrollment and received payment (ca. 12 Euros) or course credit for taking part in the experiment. The study was approved by the United Ethical Review Committee for Research in Psychology (EPKEB) in Hungary and was conducted in accordance with the Declaration of Helsinki.

**Table 1.** Descriptive data and performance on neuropsychological tests in the two groups.

| | Structured-first group<br>*n* = 25<br>*M*(*SD*) | Unstructured-first group<br>*n* = 25<br>*M*(*SD*) | Between-groups<br>difference<br>*t*/*U*/$\chi^2$ |
| --- | --- | --- | --- |
| Gender [Male/Female] | 8/17 | 8/17 | 1.00 |
| Age [years] | 21.5 (2.6) | 20.8 (1.5) | 274.50<sup>a</sup> |
| Education [years] | 14.6 (2.1) | 14.3 (1.6) | 0.69 |
| Handedness [LQ] | 53.5 (49.3) | 65.8 (28.7) | 330.50<sup>a</sup> |
| Wisconsin Card Sorting Task<br>[perseverative error percentage] | 11.67 (3.81) | 10.41 (2.02) | 1.46<sup>b</sup> |
| Corsi blocks task<br>[visuospatial short-term<br>memory span; range: 3–9] | 5.13 (0.54) | 5.04 (0.68) | 0.49 |
| Counting span task<br>[working memory span; range:<br>2–6] | 3.80 (0.86) | 4.08 (0.85) | -1.16 |
| Go/No-Go task<br>[discriminability index: hit rate<br>minus false alarm rate] | .72 (.14) | .71 (.15) | 0.40 |

*Note.* The two groups *did not differ* in any of the dependent variables and all participants performed in the
normal range on the neuropsychological tests. <sup>a</sup>In the case of violating the assumption of normality, the Mann-
Whitney U test was performed, and the *U* statistic is provided. <sup>b</sup>In case of violating the assumption of
homogeneity of variances, the robust Welch test of equality of means was performed, and the *t* statistic is
provided. Handedness was assessed with the Edinburgh Handedness Inventory revised version (Dragovic, 2004a,
2004b; Oldfield, 1971); LQ = Laterality Quotient, -100 means complete left-handedness, 100 means complete
right-handedness.

### 2.2 Experimental task

#### 2.2.1 The Alternating Serial Reaction Time (ASRT) task

Implicit acquisition of 2^nd^ order transitional probabilities was measured by a modified version of the ASRT task (J. H. Howard, Jr. & Howard, 1997; Nemeth et al., 2010; Takács et al., 2018), which was optimized for a future fMRI study using a block design. In this task, a stimulus (a dog’s head) appeared in one of four horizontally arranged empty circles on the screen (see Fig. 1C). Participants were instructed to press as quickly and accurately as possible one of the four response keys (Q, Y, M, or, O on a QWERTZ keyboard) that corresponded to the position of the stimulus (Q = leftmost position [left index finger], Y = second position from left to right [left thumb], M = third position from left to right [right thumb], O = rightmost position [right index finger]). In this task version, participants were clearly informed about the unusual mapping between spatial positions and response keys in the task instruction. During a practice phase with at least two mini-blocks of fifteen random trials each, participants had the chance to practice these stimulus-response mappings until they felt confident in proceeding to the main task. (The experimenters also required them to achieve 98% accuracy at least in the final mini block).

In the ASRT task, unbeknownst to participants, the stimuli are presented according to an eight-element sequence, within which predetermined/pattern (P) and random (r) elements alternate with one another (J. H. Howard, Jr. & Howard, 1997). For instance, 2 – r – 1 – r – 3r – 4 – r is one of the sequences, where numbers denote the four predetermined positions on the screen from left to right, and *r*s denote the randomly chosen positions out of the four possible ones (see Fig. 1A). There are 24 permutations of the four positions that could determine the applied sequence; however, because of the continuous presentation of the stimuli, there are only *six unique permutations*: 1 – r – 2 – r – 3 – r – 4 – r, 1 – r – 2 – r – 4 – r – 3 – r, 1 – r – 3 – r – 2 – r – 4 – r, 1 – r – 3 – r – 4 – r – 2 – r, 1 – r – 4 – r – 2 – r – 3 – r, and 1r – 4 – r – 3 – r – 2 – r (see also Figs. S1–S2). Note that each of these six permutations can start at any position; e.g., 1 – r – 3 – r – 4 – r – 2 – r and 2 – r – 1 – r – 3 – r – 4 – r are identical sequence permutations.

The alternating regularity yields a probability structure in which some chunks of three successive trials (triplets) occur more frequently than others. This characteristic of the task ensures that sensitivity to a biased distribution of triplets can be quantified. In the case of the 2 – r – 1 – r – 3 – r – 4 – r sequence, 2 – X – 1, 1 – X – 3, 3 – X – 4, and 4 – X – 2 triplets (X denotes the middle trial of the triplet) occur frequently since these triplets could have P – r – P or r – P – r structure. Meanwhile, for instance, 1 – X – 2 and 4 – X – 3 triplets occur less frequently since they could only have a r – P – r structure (see Fig. 1A). The former triplets are referred to as *high-probability* triplets, while the latter ones are referred to as *low-probability* triplets (e.g., Nemeth & Janacsek, 2011; Nemeth, Janacsek, Polner, & Kovacs, 2013). The construction of triplets could be considered as a method for identifying the hidden probability structure of the ASRT task. Namely, the final trial of a high-probability triplet is a probable (predictable) continuation for the first trial of the triplet, while the final trial of a low-probability triplet is a less probable continuation. For instance, in the case of the above-mentioned sequence, if the first trial of a triplet is position 3, it is more likely (with 62.5% probability) to be followed by position 4 as the third trial than either position 1, 2, or 3 (with 12.5% probability each). Each trial (stimulus) is categorized as either the third trial of a high- or a low-probability triplet. Accordingly, the construction of triplets is applied as a moving window throughout the entire stimulus set: The third trial of a triplet is also the second trial of the following triplet, and so on; thus, all stimuli are categorized this way (Kóbor, Janacsek, Takács, & Nemeth, 2017; Kóbor et al., 2018; Szegedi-Hallgató et al., 2017). There are 64 possible triplets in the task: 16 of them are high-probability triplets, and 48 are low-probability ones. With respect to the *unique* triplets, the third trials of high-probability triplets are five times more predictable based on the first trials than those of the low-probability triplets (see Figs. S1–S2).

#### 2.2.2 Generation and selection of the unstructured sequences

Besides the *structured* ASRT sequences that included the alternating regularity, *unstructured* sequences were used, in which the alternating regularity was absent. The unstructured sequences had to meet two requirements. *First*, unstructured sequences had to contain the same 64 triplets as the structured sequences; however, the probability of occurrence of each unique triplet type had to be equal. Therefore, each of the 64 triplets had to occur 30 times in any of the unstructured sequences but without the presence and repetition of the alternating regularity (1920 triplets in total, see the second requirement). In this way, unstructured sequences could also be considered as pseudorandom sequences with the constraint that all triplets occurred with equal probability (25%). *Second*, unstructured sequences had to contain the same number of trials as structured sequences because they determined stimulus presentation in an equal number of blocks. This meant the presentation of altogether 1920 triplets distributed over 48 blocks with 40 triplets in each, respectively (see below). For this purpose, *without* the use of the alternating regularity, several trial sets were generated in MATLAB 2015a (The MathWorks Inc., Natick, 224 MA, USA). Particularly, by randomly changing one trial of the trial sets at a time, the trial set minimizing the deviation from the optimal 30 times of occurrence was selected (the maximal error was set to two). Using this algorithm, a dozen trial sets satisfying this criterion were kept.

These trial sets were then subjected to three further constraints: (1) the maximal repetition of a unique triplet in any of the blocks could be no more than four; (2) the maximal immediate repetition of a trial [position] could be no more than five across the entire trial set; (3) in larger time bins (16 blocks) of the unstructured trial set, the overall occurrence probability of triplets that can be categorized as high- vs. low-probability in the structured ASRT sequences should approximate 25% and 75%, respectively, since there are 16 unique high-probability and 48 unique low-probability triplets for a given ASRT sequence (see above the ASRT task description). This third constraint ensured that at the level of unique triplets, the transitional probabilities were equal. Six of the trial sets were appropriate regarding constraints (1) and (2). Stimuli of these six trial sets were categorized into *triplets* following either of the six unique structured ASRT sequences (see Fig. 1B); and, constraint (3), i.e., the ratio of the high- and low-probability triplets, was checked on these categorized trial sets. Finally, altogether 19 trial sets satisfied all three constraints and were kept using as *unstructured sequences*.

When assigning the structured ASRT and unstructured sequences to participants, we ensured that the distribution of the six unique ASRT sequence types was even across the two groups. The 1 – r – 2 – r – 3 – r – 4 – r sequence was used five times, and all the other sequences were used four times in each of the groups (i.e., for 25 participants per group). For each participant, the selection of a sequence from the six unique types was pseudorandom. The applied files containing the structured ASRT and unstructured sequences were matched one-to-one across the two groups. Note that for each respective participant, at the level of unique triplets, the identified high- and low-probability triplets were the same in the unstructured sequences as in the structured ASRT sequences (see Fig. 1A–B). Indeed, the probability of triplets differed only in the structured sequences and their probability was equal in the unstructured sequences (see Figs. S1–S2). Meanwhile, in the remainder of the paper, triplets in the unstructured sequences are still referred to as either high- or low-probability according to their actual probability in the structured sequences.

As a result of the procedure used for generating, selecting, and matching the sequences, in the present sample, the distribution of high- and low-probability triplets did not differ across the four stimulus positions either in the structured ASRT (χ^2^(3) = 4.86, *p* = .183) or in the unstructured sequences (χ^2^(3) = 0.02, *p* = .999); in addition, these associations between triplet distribution and stimulus position did not differ across the sequence types (Wald χ^2^(3) = 2.34, *p* = .504). When the high- and low-probability triplet categories were collapsed, the distribution of stimulus positions across sequence types also did not differ (χ^2^(3) = 1.56, *p* = .670). In this way, lower-level characteristics of the sequences would not account for the assumed between-sequence RT variations related to acquiring the 2^nd^ order transitional probability structure (cf. Reed & Johnson, 1994).

### 2.3 Procedure

An experimental trial started with the presentation of the stimulus at one of the four positions for 500 ms. After stimulus offset, the image of the four positions was displayed for 200 ms. Then, the next trial started, yielding a 700-ms-long inter-trial interval. The behavioral response (keypress) was expected during the whole trial from stimulus onset until the end of the trial (i.e., for altogether 700 ms, see Fig. 1C). These trial events were always the same with fixed durations, irrespective of whether participants provided correct, incorrect, or missing response(s). In this task version, no feedback was presented as a function of the quality of the response. The lack of feedback presentation and the fact that participants could proceed with the trial without providing the correct response ensured that each trial and each block had the same lengths, respectively. Importantly, only correctly responded trials were analyzed in the present study.

One block of the task contained 42 trials. There were 48 blocks with the structured ASRT sequence and 48 blocks with the unstructured sequence. In each of the structured blocks, the eight-element-long alternating regularity repeated five times after two starter trials that were not categorized as triplet elements (since also the foremost triplet technically required three successive trials). The alternating regularity was missing from the unstructured blocks, but, as in the structured blocks, 40 triplets followed the two starter trials that were not categorized as triplet elements. After each block, participants received feedback about their mean reaction time and accuracy in the given block. The length of this between-blocks “rest period” with feedback was jittered to be methodologically optimal for a future fMRI experiment (it lasted for 10, 12, or 14 sec [mean = 12 sec]). Altogether 96 blocks were completed (4032 trials in total).

The Structured-first group completed 48 structured blocks followed by 48 unstructured blocks. The Unstructured-first group completed 48 unstructured blocks followed by 48 structured blocks. All participants proceeded with the task from its structured/unstructured to unstructured/structured half without receiving information about any change in the task (see Fig. 1D). Two breaks (1.5 mins each) were inserted after the 32^nd^ and 64^th^ blocks, in which participants could have had a short rest. The experimental procedure lasted about 1.5 hours including the administration of a short post-task questionnaire. This assessed participants’ task-solving strategies and their consciously accessible knowledge about the structure of the task and the transitional probabilities (Kóbor et al., 2017; Nemeth, Janacsek, & Fiser, 2013; Song, Howard, & Howard, 2007). Namely, participants were asked whether (1) they followed any task-solving strategies to improve their performance, (2) if yes, to what extent they found it efficient; (3) whether they noticed anything special regarding the task; (4) whether they noticed any regularity in the sequence of stimuli; and (5) whether they noticed any substantial change in the sequence of stimuli. The first author (AK) qualitatively rated participants’ answers to questions (1) and (2), and rated the answers to questions (3), (4), and (5) on a 5-item scale (1 = “Nothing noticed”, 5 = “Total awareness”). None of the participants reliably reported noticing the alternating regularity, the presence and repetitions of the triplets, or any change in the stimulus sequence between the task halves (the mean score for the three questions was 1.006, *SD* = 0.082). Although participants reported several strategies they found somewhat facilitating (e.g., counting the stimuli, fixating to the center of the screen, catching the rhythm of trials by silently singing, bouncing their legs, or moving their fingers), these were unrelated to the hidden structure of the task. Only one participant reported trying to search for some “logic” in the sequence but as a subjectively inefficient strategy.

The current ASRT task version was written in and controlled by MATLAB 2015a using the Psychophysics Toolbox Version 3 (PTB-3) extensions (Brainard, 1997; Pelli, 1997). Stimuli were displayed on a 15” LCD screen at a viewing distance of 100 cm. Neuropsychological tests (see Participants section) were administered a few days before the main experiment during a one-hour-long session.

### 2.4 Statistical analysis

Following the standard data analysis protocol established in previous studies using the ASRT task (e.g., J. H. Howard, Jr. & Howard, 1997; Kóbor et al., 2017; Nemeth, Janacsek, Polner, et al., 2013; Song et al., 2007; Virag et al., 2015), two types of low-probability triplets – repetitions (e.g., 1 – 1 – 1, 4 – 4 – 4) and trills (e.g., 1 – 2 – 1, 2 – 4 – 2, see Fig. 1B) – were eliminated from the analyses because preexisting response tendencies have often been shown to them (D. V. Howard et al., 2004). In addition, eight-block-long units of the task were collapsed into larger time bins labeled as *epochs*, yielding altogether six structured epochs (containing the ASRT sequence) and six unstructured epochs (containing the unstructured sequence). From this point of view, while the Structured-first group performed six structured epochs followed by six unstructured epochs, the Unstructured-first group performed six unstructured epochs followed by six structured epochs. Epochs are labeled consecutively in this paper (1, 2, etc.) within each sequence type. For each participant and epoch, separately for high- and low-probability triplets, median RT was calculated only for correct responses.

Triplet learning on the RTs, i.e., faster RTs to high-probability than to low-probability triplets, was first quantified with a four-way mixed design analysis of variance (ANOVA) with Sequence (structured vs. unstructured), Triplet (high-vs. low-probability), and Epoch (1–6) as within-subjects factors and Group (Structured-first group vs. Unstructured-first group) as a between-subjects factor. Second, to more directly test the change in triplet learning as a function of the different sequence types, three-way mixed ANOVAs with Triplet and Epoch as within-subjects factors and Group as a between-subjects factor were performed on the RTs related separately to the structured and unstructured epochs. In all ANOVAs, the Greenhouse-Geisser epsilon (ε) correction (Greenhouse & Geisser, 1959) was used when necessary. Original *df* values and corrected (if applicable) *p* values are reported together with partial eta-squared (η_*p*_^2^) as the measure of effect size. LSD (Least Significant Difference) tests for pairwise comparisons were used to control for Type I error.

Regarding the possible experimental effects and their interpretation, the Triplet main effect implies *triplet learning* (faster RTs to high- than to low-probability triplets) and the Triplet * Epoch interaction implies *changes in triplet learning* as the task progresses, usually an increase across epochs (e.g., Janacsek, Ambrus, Paulus, Antal, & Nemeth, 2015; Kóbor et al., 2017; Nemeth et al., 2010; Nemeth, Janacsek, Polner, et al., 2013; Takács et al., 2017; Tóth et al., 2017). The Epoch main effect implies *general skill (RT) improvements* reflecting more efficient visuomotor and motor-motor coordination due to practice (Hallgató et al., 2013; Juhasz, Nemeth, & Janacsek, 2019). The *prior knowledge effect* is indicated by the Sequence * Triplet * Group and/or the Sequence * Triplet * Epoch * Group interactions: Namely, if prior knowledge of the transitional probabilities influences later stimulus processing, triplet learning per se or its change over time should differ between structured and unstructured epochs *and* across the two groups experiencing the structured and unstructured epochs in the opposite order. In the Results section below, we use these terms when describing the observed statistical effects.

To follow up the prior knowledge effect, triplet learning scores in the structured and unstructured epochs were calculated as the RT difference between the triplet types (RTs to low-probability triplets minus RTs to high-probability triplets). *Overall triplet learning scores* were considered for the structured and unstructured epochs, respectively, as the mean of the scores calculated for each of the six epochs. The overall triplet learning scores for the structured and unstructured epochs were first compared within each group. Then, these scores were compared between the groups. Finally, the change in mean RTs across the structured and unstructured epochs separately for the high- and low-probability triplets were compared within each group.

To test the *persistence* of the prior knowledge effect, triplet learning scores were further analyzed in the Structured-first group. To find a balance between increased power and capturing the time course of persistence, triplet learning scores were averaged over *two consecutive epochs* (i.e., “thirds”) of the structured and unstructured sequences, respectively. Then, these scores were compared against zero in each sequence type. Finally, these scores were compared between the corresponding thirds of the structured and unstructured sequences.

## 3. Results

### 3.1 Overall analysis

The Sequence (structured vs. unstructured) by Triplet (high-vs. low-probability) by Epoch (1–6) by Group (Structured-first group vs. Unstructured-first group) *overall* ANOVA on the RTs revealed the significant main effects of Sequence, *F*(1, 48) = 6.77, *p* = .012, η_*p*_^2^ = .124, Triplet, *F*(1, 48) = 45.90, *p* < .001, η_*p*_^2^ = .489, and Epoch, *F*(5, 240) = 35.91, ε = .612, *p* < .001, η_*p*_^2^ = .428. As these main effects were qualified by significant higher-order interactions, only the latter effects are detailed below.

#### 3.1.1 Triplet learning

The Sequence * Triplet, *F*(1, 48) = 22.92, *p* < .001, η_*p*_^2^ = .323, and the Triplet * Epoch, *F*(5, 240) = 2.42, *p* = .036, η_*p*_^2^ = .048, interactions were significant, while the Sequence * Triplet * Epoch, *F*(5, 240) = 2.18, ε = .814, *p* = .072, η_*p*_^2^ = .043 interaction was a trend level. These effects indicated that the change in triplet learning over the course of the task differed between the structured and unstructured epochs.

#### 3.1.2 General skill improvements

The significant Sequence * Group, *F*(1, 48) = 69.87, *p* < .001, η_*p*_^2^ = .593, and the Sequence * Epoch * Group, *F*(5, 240) = 33.59, ε = .706, *p* < .001, η_*p*_^2^ = .412, interactions showed that between-groups differences emerged as a function of first experiencing the structured or the unstructured half (i.e., six epochs) of the task. Particularly, while the Structured-first group became increasingly faster over the structured epochs due to practice and showed similar RTs over the unstructured epochs, this was reversed in the Unstructured-first group, where increasingly faster RTs were observed over the unstructured epochs and similar RTs over the structured epochs (see Fig. 2). This effect suggests that general skill improvements were found in the first half of the task, irrespective of whether this half was structured or unstructured and the distribution of triplets. Relatedly, the following *nonsignificant* main effects and interactions emerged: main effect of Group, *F*(1, 48) = 1.96, *p* = .168, η_*p*_^2^ = .039, Epoch * Group interaction, *F*(5, 240) = 0.47, ε = .612, *p* = .706, η_*p*_^2^ = .010, and Sequence * Epoch interaction, *F*(5, 240) = 1.20, ε = .706, *p* = .313, η_*p*_^2^ = .024.

**Figure 2.**
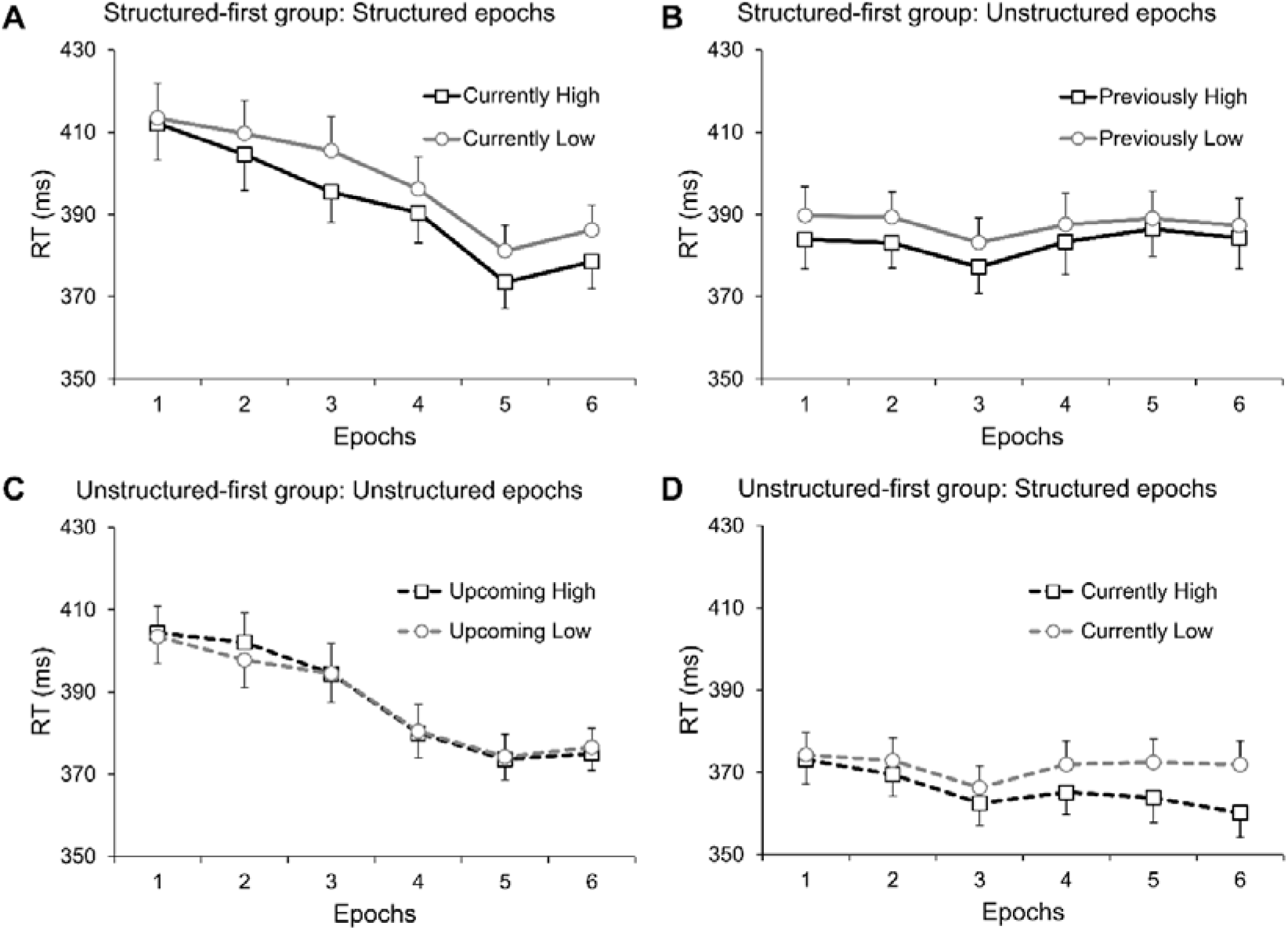
Temporal dynamics of triplet learning across groups and sequence types. Group-average RTs (A– B: Structured-first group; C–D: Unstructured-first group) for correct responses as a function of time bin (epochs 1–6) and triplet type (currently/previously/upcoming high-vs. low-probability triplets, according to their actual probability in the given sequence and the order of the sequence in the given group) are presented in the structured (A, D) and unstructured (B, C) epochs. Error bars denote standard error of mean.

#### 3.1.3 Prior knowledge effect

The significant Triplet * Group interaction, *F*(1, 48) = 4.92, *p* = .031, η_*p*_^2^ = .093, was qualified by the significant Sequence * Triplet * Group interaction, *F*(1, 48) = 7.96, *p* = .007, η_*p*_^2^ = .142. Importantly, the latter indicated that the difference in triplet learning between the structured and unstructured epochs varied across the groups, which is regarded as the prior knowledge effect (see Fig. 2). This effect did not reliably vary as a function of practice with the task, as shown by the nonsignificant Sequence * Triplet * Epoch * Group interaction, *F*(5, 240) = 0.61, ε = .814, *p* = .661, η_*p*_^2^ = .012. Relatedly, the modulating effect of structured vs. unstructured epochs on triplet learning was also supported by the significant Triplet * Epoch * Group interaction, *F*(5, 240) = 2.31, *p* = .045, η_*p*_^2^ = .046, showing that if both task halves with structured and unstructured epochs were collapsed, the trajectory of triplet learning would differ across the groups.

### 3.2 Follow-up of the prior knowledge effect

To follow up the prior knowledge effect, pairwise comparisons contrasting the overall triplet learning scores were performed (see Fig. 3). In the Structured-first group, the triplet learning score was similar between the structured and unstructured epochs (6.3 ms vs. 4.6 ms, *p* = .171). Thus, the behavioral effect of biased triplet probabilities (i.e., high- and low-probability triplets in the structured epochs) persisted even after this bias was eliminated (i.e., the unique triplets occurred with equal probability in the unstructured epochs). Meanwhile, in the Unstructured-first group, the triplet learning score was significantly higher over the structured epochs than over the unstructured epochs; in the latter, it was virtually zero (5.9 ms vs. −0.4 ms, *p* < .001). In addition, the triplet learning score did not differ between the groups over the structured epochs (6.3 ms vs. 5.9 ms, *p* = .840), but it was significantly higher in the Structured-first group than in the Unstructured-first group over the unstructured epochs (4.6 ms vs. −0.4 ms, *p* < .001).

**Figure 3.**
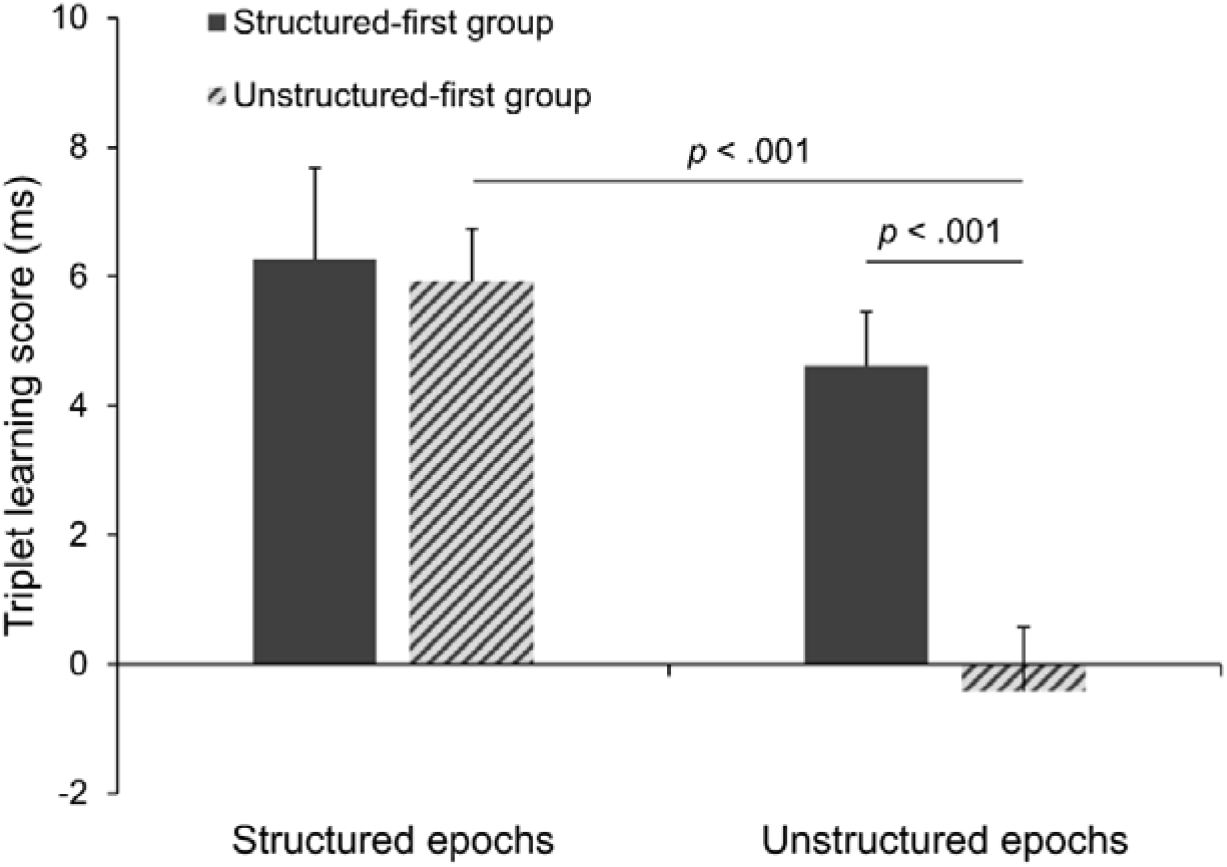
Persistence of the acquired implicit knowledge. Group-average overall (mean) triplet learning scores (RTs to low- minus RTs to high-probability triplets) are presented in the Structured-first and Unstructured-first groups for the structured and unstructured epochs, respectively. Error bars denote standard error of mean.

In the Structured-first group, mean RTs on the high- and low-probability triplets decreased from the structured to the unstructured epochs to a similar extent (high-probability triplets: 392 ms to 383 ms [Diff = 9 ms], *p* = .003; low-probability triplets: 399 ms to 388 ms [Diff = 11 ms], *p* < .001; the difference in RT decrease between high- and low-probability triplets was not significant, *p* = .226), indicating only general skill improvements from the structured to the unstructured epochs. In contrast, in the Unstructured-first group, a larger RT decrease was found on the high-probability triplets (388 ms to 366 ms [Diff = 22 ms], *p* < .001) than on the low-probability ones (388 ms to 372 ms [Diff = 16 ms], *p* < .001; the difference in RT decrease between high- and low-probability triplets was significant, *p* < .001) from the unstructured to the structured epochs, indicating triplet learning over the structured epochs.

### 3.3 Persistence of the prior knowledge effect

To test the persistence of the prior knowledge effect in the Structured-first group, triplet learning scores were averaged over epoch_1_ and epoch_2_ (M_e1e2_), epoch_3_ and epoch_4_ (M_e3e4_), epoch_5_ and epoch_6_ (M_e5e6_), respectively. These new scores were compared against zero and between the structured and unstructured sequences. The obtained results are presented in Figure 4A and detailed below.

**Figure 4.**
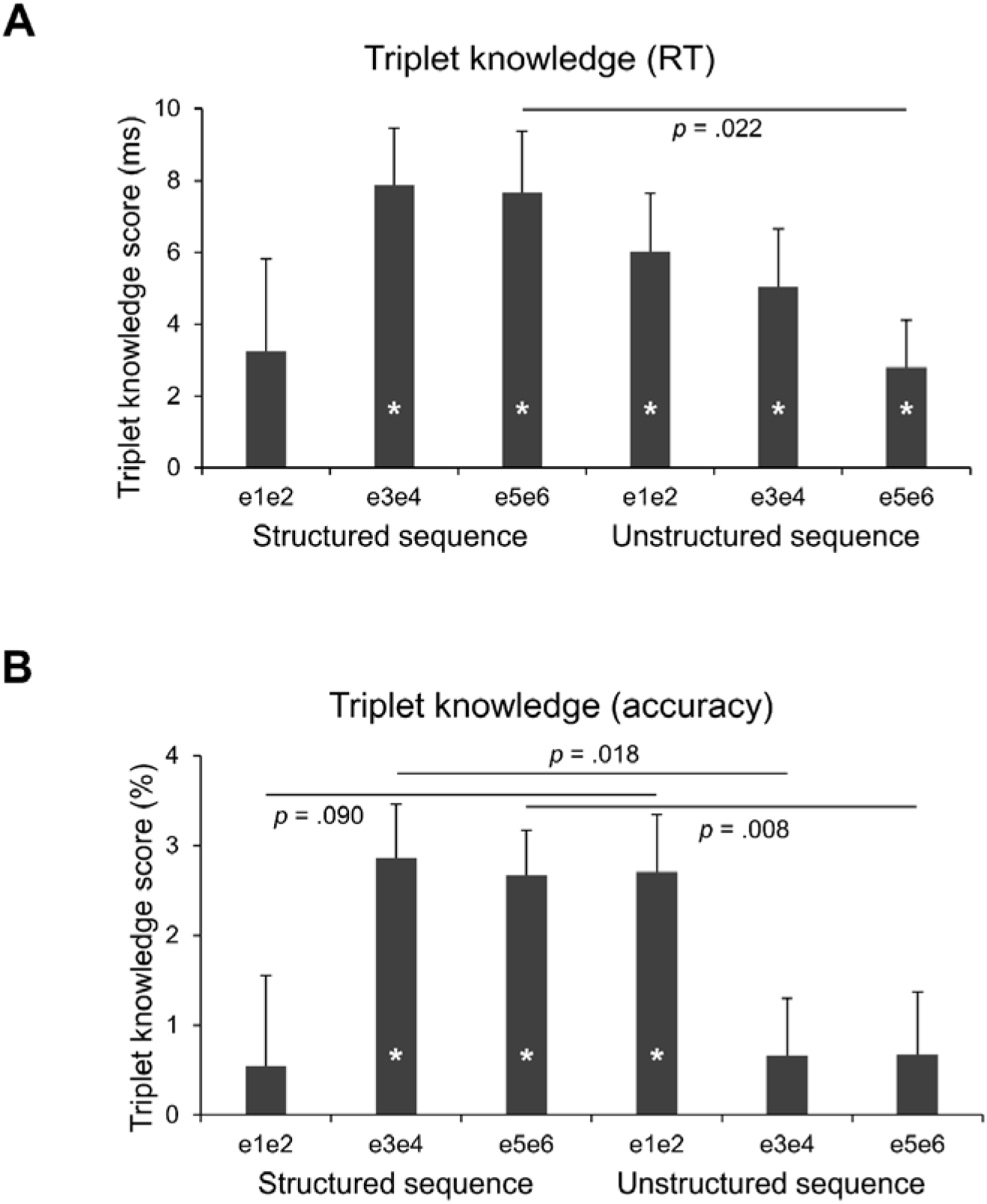
Temporal characteristics of persistency. In the Structured-first group, group-average triplet knowledge scores measured by RTs (A) and accuracy (B) averaged over two consecutive epochs (i.e., “thirds”) of the structured and unstructured sequences, respectively, are presented. Error bars denote standard error of mean. White asterisks denote that the given score significantly differs from zero (*p* < .050).

The extent of triplet knowledge differed significantly from zero over all thirds of the structured and unstructured sequences (all *t*s ≥ 2.11, *p*s ≤ .045), except for the very first one at the beginning of the task (*t*(24) = 1.26, *p* = .221). This indicated the presence of triplet knowledge from the second third of the structured sequence throughout the task. Triplet knowledge did not differ between the unstructured and structured sequences during the first (M_e1e2_: 6.0 ms vs. 3.2 ms, respectively, *t*(24) = −0.96, *p* = .349) and second thirds of the task (M_e3e4_: 5.1 ms vs. 7.9 ms, respectively, *t*(24) = 1.35, *p* = .190). In contrast, triplet knowledge over the last third of the unstructured sequence was significantly lower than over the last third of the structured sequence (M_e5e6_: 2.8 ms vs. 7.7 ms, respectively, *t*(24) = 2.46, *p* = .022). The decreasing extent of triplet knowledge is also noticeable in Figure 2B as RTs to high- and low-probability triplets approaching one another in the last third of the unstructured sequence. These results altogether suggest that participants acquired the triplet knowledge in the second third of the structured sequence; and, after biased probabilities had been removed from the stimulus stream, the update of the triplet knowledge was evident in behavior only in the final third of the unstructured sequence.

### 3.4 Separate analysis of the structured and unstructured epochs

The Triplet by Epoch by Group ANOVA on the RTs related to the *structured* epochs revealed the significant main effects of Triplet, *F*(1, 48) = 56.00, *p* < .001, η_*p*_^2^ = .538, Epoch, *F*(5, 240) = 27.54, ε = .700, *p* < .001, η_*p*_^2^ = .365, and Group, *F*(1, 48) = 9.25, *p* = .004, η_*p*_^2^ = .162. These effects were qualified by the significant Triplet * Epoch, *F*(5, 240) = 4.55, *p* = .001, η_*p*_^2^ = .087, and Epoch * Group, *F*(5, 240) = 15.77, ε = .700, *p* < .001, η_*p*_^2^ = .247, interactions, indicating that triplet learning increased with practice and general skill improvements differed between the groups (see Fig. 2A, D). Triplet learning and its change over the structured epochs did not differ between the groups, as shown by the nonsignificant Triplet * Group, *F*(1, 48) = 0.04, *p* = .840, η_*p*_^2^ = .001, and Triplet * Epoch * Group interactions, *F*(5, 240) = 1.54, *p* = .177, η_*p*_^2^ = .031.

The same Triplet by Epoch by Group ANOVA on the RTs related to the *unstructured* epochs revealed the significant main effects of Triplet, *F*(1, 48) = 10.61, *p* = .002, η_*p*_^2^ = .181, and Epoch, *F*(5, 240) = 13.45, ε = .605, *p* < .001, η_*p*_^2^ = .219, while the Triplet * Epoch interaction was not significant, *F*(5, 240) = 0.24, *p* = .945, η_*p*_^2^ = .005). Importantly, the significant Triplet * Group interaction, *F*(1, 48) = 15.23, *p* < .001, η_*p*_^2^ = .241, showed that triplet learning was larger in the Structured-first group than in the Unstructured-first group (4.6 ms vs. −0.4 ms) over the unstructured epochs, but this did not change with time (nonsignificant Triplet * Epoch * Group interaction, *F*(5, 240) = 1.33, *p* = .251, η_*p*_^2^ = .027). The trajectory of general skill improvements differed between the groups (significant Epoch * Group interaction, *F*(5, 240) = 16.12, ε = .605, *p* < .001, η_*p*_^2^ = .251, see Fig. 2B, C), and the groups did not differ in overall RT (nonsignificant Group main effect, *F*(1, 48) = 0.09, *p* = .764, η_*p*_^2^ = .002) over the unstructured epochs.

### 3.5 Analysis of accuracy

Since only the RTs of the correctly responded trials were analyzed, it should be ensured that the two groups did not differ in accuracy. Therefore, the Sequence by Triplet by Epoch by Group ANOVA was also conducted on accuracy data (calculated as the ratio of correct responses for each participant and epoch, separately for high- and low-probability triplets). As indicated by the nonsignificant main effect of Group, *F*(1, 48) = 1.53, *p* = .221, η_*p*_^2^ = .031, the two groups were comparable in overall accuracy (Structured-first group: 86.4%; Unstructured-first group: 88.1%).

Although the analysis of accuracy is not the focus of this study, for the sake of completeness, we provide the other significant main effects and interactions revealed in this ANOVA, but these are not detailed. The main effects of Triplet, *F*(1, 48) = 25.25, *p* < .001, η_*p*_^2^ = .345, and Epoch, *F*(5, 240) = 7.42, ε = .639, *p* < .001, η_*p*_^2^ = .134, were significant. Relatedly, the Sequence * Triplet, *F*(1, 48) = 4.91, *p* = .032, η_*p*_^2^ = .093, and the Sequence * Triplet * Epoch, *F*(5, 240) = 3.49, *p* = .005, η_*p*_^2^ = .068, interactions were also significant. In brief, responses to high-probability triplets were more accurate than those to low-probability ones. However, while this difference in accuracy increased over the structured epochs, it gradually decreased over the unstructured epochs.

The Sequence * Group, *F*(1, 48) = 9.75, *p* = .003, η_*p*_^2^ = .169, and the Sequence * Epoch * Group, *F*(5, 240) = 7.44, ε = .524, *p* < .001, η_*p*_^2^ = .134, interactions were significant, as well. The Sequence * Triplet * Epoch * Group interaction was a trend level, *F*(5, 240) = 1.92, *p* = .091, η_*p*_^2^ = .039. The latter effect indicated that the above-described Sequence * Triplet * Epoch interaction was mostly driven by the responding pattern of the Structured-first group. Particularly, in the Structured-first group, while the difference in accuracy between high- and low-probability triplets increased over the structured epochs, it tended to decrease over the unstructured epochs. In the Unstructured-first group, accuracy between high- and low-probability triplets differed only over the structured epochs.

The results of comparing triplet knowledge measured by accuracy calculated for each third of each sequence in the Structured-first group were in line with these effects (see Fig. 4B). The extent of triplet knowledge differed significantly from zero over the middle and last thirds of the structured sequence and over the first third of the unstructured sequence (all *t*s ≥ 4.24, *p*s < .001). Accordingly, triplet knowledge tended to be higher over the first third of the unstructured sequence than over the first third of the structured sequence (M_e1e2_: 2.7% vs. 0.6%, respectively, *t*(24) = −1.77, *p* = .090). In contrast, triplet knowledge was significantly lower over the last two thirds of the unstructured sequence than over the corresponding thirds of the structured sequence (M_e3e4_: 0.7% vs. 2.9%, respectively, *t*(24) = 2.55, *p* = .018; M_e5e6_: 0.7% vs. 2.7%, respectively, *t*(24) = 2.91, *p* = .008). These results suggest that updating the triplet knowledge after biased probabilities had been removed was as fast as acquiring that knowledge in the first place.

## 4. Discussion

### 4.1 Summary of results

This study investigated whether the implicitly acquired knowledge of a 2^nd^ order transitional probability structure influenced the processing of unpredictable transitional probabilities across phases of a learning task. To this end, the changes in RTs to more probable and less probable short-range transitional probabilities (triplets) embedded in a stimulus sequence were tracked. The stimulus sequence changed over the experimental task because biased triplet probabilities were present in one-half of the task blocks and absent in the other half, without explicitly denoting this change at the surface level.

In line with our assumptions, while the participant group completing the structured half of the task first showed triplet learning across both the structured and unstructured blocks, the participant group completing the unstructured half first showed triplet learning only over the structured blocks and not over the unstructured blocks. Based on the performance of the group completing the structured half of the task first, it seems that the already acquired implicit knowledge of the short-range transitional probabilities persisted across the learning phases, even after the bias in the transitional probabilities had been removed. This persistence characterized two-thirds of the unstructured blocks. Then, the update of prior knowledge became evident in RT triplet learning over the last third of these blocks. The results also imply that the updating process took longer than the primary acquisition, which required only one-third of the structured blocks. Based on the performance of the group completing the unstructured half of the task first, it seems that the protracted exposure to unbiased transitional probabilities did not negatively influence the acquisition of the biased transitional probabilities later in the task. Therefore, any potential expectation or knowledge built upon the pseudorandom stimuli was also updated to promote the acquisition of the newly experienced biased transitional probabilities.

### 4.2 Persistent implicit statistical knowledge

Participants completing the structured blocks first similarly perceived the relations of stimuli in both types of blocks. That is, their perception could have been influenced by the primarily experienced transitional probability structure. In other words, because of the task environment, they might have worked up a tendency towards pattern detection, which could have resulted in forming implicit expectations about the upcoming stimuli. Then, these expectations remained persistent throughout the task.

To explain the observed persistency, one should consider the characteristics of the given learning environment. In the present task, acquisition happened in an incidental and implicit manner: our participants did not know that they were in a learning situation, they did not have information on whether the sequence of stimuli was random or followed any underlying pattern, the critical change point between the task halves remained unnoticed, and they were not required to actively or explicitly predict the probability of the next stimulus. In addition, they did not receive feedback (or any reward) on the correctness of their responses. Meanwhile, participants were instructed to maintain a certain level of speed and accuracy, but, for them, this was the only explicit goal of the task. They successfully achieved this goal, as shown by the behavioral results indicating general skill improvements due to practice. This performance improvement was mostly independent of the change in the underlying probability structure. Therefore, it seems that as participants had gained confidence in task solving, they followed their already established, automatized strategy on stimulus processing and responding (cf. Karuza et al., 2016). As the surface of the task remained consistent, they had no reason to doubt their current implicit beliefs about the probability of the upcoming stimulus (cf. Zinszer & Weiss, 2013). This might have promoted the persistently faster processing of the stimuli that occurred with high probability only in the previous task half.

From a broader perspective, it could be adaptive that the acquired representations of the structured stimuli remain persistent over time and robust to change. This way, the representations could remain sensitive to the primary transitional probability structure later in time, although other structures are simultaneously acquired (cf. Gebhart et al., 2009; Qian et al., 2012; Todd et al., 2020; Todd et al., 2011). This might be even more pronounced if no explicit cue or performance deterioration signals the need for an updating process. Supporting this notion, during the acquisition of two different statistical structures, a tendency towards neural efficiency coupled with diminished sampling of the input underlying the second statistical structure has been shown (Karuza et al., 2016). Accordingly, we assume that such a “processing efficiency” might explain our results on the transition from structured to pseudorandom stimuli.

### 4.3 Temporal characteristics of persistency

It has already been demonstrated that the primarily acquired implicit knowledge of the biased distribution of transitional probabilities remains stable over longer time periods, such as one week (Nemeth & Janacsek, 2011) or even one year (Romano, Howard, & Howard, 2010). Moreover, this knowledge is resistant to short periods of interfering sequences (that partially overlap with the primarily practiced sequence) not only after 24 hours but also after one year (Kóbor et al., 2017). The present study could extend these results on persistency as follows. If there is an essential change in the stimulus probabilities characterizing the given environment, the duration required for update the existing probabilistic representations seems to be longer than the duration required for acquiring these representations, at least at the behavioral level.

In detail, for participants completing the structured blocks first, acquiring the biased probability structure required one-third of the structured sequence; then, this knowledge remained persistent until the end of the task. However, participants were also sensitive to the lack of bias or the altered probability of the unique triplets over the unstructured sequence. Particularly, while triplet knowledge measured by RTs was comparable over the first two-thirds of the structured vs. unstructured sequences, triplet knowledge was decreased in the last third of the unstructured sequence as compared with the structured sequence. Thus, participants needed to complete two-thirds of the unstructured sequence to update their existing knowledge of the probability structure, which was acquired after the completion of only one-third of the structured sequence. When triplet knowledge was defined by differences in accuracy, updating was more pronounced and became evident earlier: Triplet knowledge was lower over the middle and last thirds of unstructured sequence than that of the structured sequence. Indeed, after one-third of the task blocks, triplet knowledge abruptly increased when the bias was present and dropped when the bias was eliminated. Overall, the results of participants completing the structured blocks first might indicate an “implicit need” for updating the prior representations of the probability structure. At the same time, these results also highlight the constraining effect of the primarily acquired, possibly overlearned statistical structure in the adaptation to a new environment (cf. Bulgarelli & Weiss, 2016; Gebhart et al., 2009).

It has been suggested that successful adaptation to a range of similar tasks requires the forgetting or weakening of some specific features of the already acquired representations (Robertson, 2018). As described above, in the present case, this would have been the forgetting of the initial triplet probability information when starting the other task half. It is plausible to assume that after having had performed even more unstructured blocks, participants would have learned that the previously high-probability triplets no longer occurred with higher probability than the previously low-probability ones, and, therefore, the initial triplet probability information would have been forgotten or “unlearned”. However, in accordance with the findings of Szegedi-Hallgató et al. (2017) indicating the coexistence of the previously and the recently acquired implicit knowledge of the changed statistical structure in the ASRT task, it is more likely that, at the level of triplet representations, no “extinction” or “unlearning” happened. Evidence from human and animal studies suggests that adaptation to different contexts does not involve the complete removal of representations of the previous context (Bulgarelli & Weiss, 2016; Gordon, Bilolikar, Hodhod, & Thomas, 2020; Qian et al., 2012). Instead, the formation of new representations, the reconsolidation or inhibition of the previously created ones, and the switching between multiple representations seem more likely (Chandler & Gass, 2013). Either of the latter processes supports the interpretation that in the present experiment, stimulus processing was determined by prior representations of the transitional probability structure that changed slowly with accumulating experiences about the ongoing stimulus context (cf. Daw et al., 2011; Griffiths et al., 2008; Shohamy & Daw, 2015). The exact mechanisms by which this slow change might have occurred has yet to be determined, and formal models should be developed to investigate the temporal dynamics of these mechanisms (cf. R. Frost et al., 2019; Karuza et al., 2016; Qian et al., 2012; Zhao et al., 2019).

### 4.4 Exposure to pseudorandom stimuli

Over the structured blocks, participants completing the unstructured blocks first showed a triplet learning trajectory comparable to that of the other group (nonsignificant Triplet * Group and Triplet * Epoch * Group interactions, see also Fig. 2A, D). With an unsigned change in the probability structure, the pervasive experience with pseudorandom stimuli could have also caused an entrenchment effect. In this case, these participants could have started the acquisition of the biased probability structure with some disadvantage or could have showed the complete lack of triplet learning when exposed to the structured blocks. Instead, according to the results, it seems that prior experience with equal transitional probabilities did not negatively influence the further acquisition of the biased probability structure.

As an explanation for the triplet learning performance of these participants, it is conceivable that they might have primarily established a wider hypothesis space about the properties of the stimuli, since they could not extract complex transitional probabilities over the unstructured blocks. Such representations could be useful if the stimuli considered as random in the given environment with limited observations in fact followed some structure. With a wider hypothesis space, stimulus processing and acquisition might have proceeded flexibly, enabling the acquisition of the statistical structure when it was indeed present. In support of this idea, similar results were found in a binary choice task testing probability-matching behavior (i.e., matching choice probabilities to outcome probabilities instead of the optimal maximizing strategy). In that task, the transition between the no pattern (no serial dependence in the sequence) and pattern half (repeating deterministic sequence) was clearly indicated. Results showed that participants who were more prone to search for patterns in the no pattern half of the task showed higher accuracy in the pattern half as compared with those participants who were less prone to follow any complex search strategy (Gaissmaier & Schooler, 2008).

### 4.5 Perception and acquisition of changing statistical structures

Earlier studies using different paradigms showed mixed results on how individuals updated their already acquired knowledge of the underlying probabilities when these probabilities changed. It was found previously that individuals accurately estimated the hidden probability parameter of a nonstationary stochastic environment as well as quickly updated their estimates (Gallistel, Krishan, Liu, Miller, & Latham, 2014). In this task, participants assessed the proportion of one stimulus category and had the opportunity to update their estimates on a trial-by-trial basis. Importantly, at the beginning of the task, they were told that probabilities could unexpectedly change. Similarly, in another experiment where subsequent numerical values had to be predicted, participants updated each prediction as a function of their explicitly denoted prediction errors. By tracking the prediction errors, after an unsigned change in the distribution of the values during the task, participants could adjust their predictions (Nassar, Wilson, Heasly, & Gold, 2010). The quick adaptation to changing probabilities was also observed when choosing between two options associated with different probabilities and reward magnitudes, i.e., with clear feedback signals (Behrens, Woolrich, Walton, & Rushworth, 2007).

These studies altogether suggest that effective decision making necessitates the continuous tracking of the environmental probabilities and the evaluation of each signal that possibly implies a change in these probabilities. However, these observations have been derived from situations in which active agents were required to make explicit decisions that pertained directly to the probabilistic features of the ongoing task modeling volatile environments. The latter characteristics possibly explain why the present results contrast with earlier findings. The ASRT task used in this study should not be considered as a(n) (explicit) decision-making or probabilistic reinforcement-learning paradigm in which quick updating of beliefs could happen (Bulgarelli & Weiss, 2016). Instead, the type of learning that this task measures more likely fits into the category of unsupervised statistical learning (Fiser & Aslin, 2001). It intends to model a stable stimulus environment with low volatility where hidden probabilistic regularities occur interspersed with noise in the form of nonadjacent transitional probabilities. These features could contribute to the persistence of the acquired regularities rather than to the abrupt change of the related representations.

A paradigm more similar to the ASRT task is the classical serial reaction time (SRT) task (e.g., Nissen & Bullemer, 1987), where a repeating deterministic sequence guides stimulus presentation in the structured blocks. In the SRT, performance usually deteriorates on the unstructured blocks with random or pseudorandom stimuli, meaning that RTs suddenly increase compared to the level reached by the end of the last structured block. Meanwhile, in the present study, participants completing the structured blocks first showed persistent triplet learning performance over several blocks of pseudorandom stimuli in terms of RTs. It is possible that in the case of probabilistic sequences (used in the ASRT task) as opposed to deterministic ones, the acquisition processes are more sensitive to smooth transitions between stimuli or chunks of stimuli. This specific sensitivity evolved in participants practicing the structured blocks first might have led the extraction of triplets even over the unstructured blocks of the task (see also the Transfer of prior knowledge section). In addition, as opposed to deterministic and pseudorandom sequences, the probabilistic sequence might have provided learnable but sufficiently novel information on a trial-by-trial basis (Maheu, Meyniel, & Dehaene, 2020), promoting the relatively fast acquisition of the statistical structure as well as its persistence.

Other studies using linguistic stimuli with transitional probabilities have started to investigate how to attenuate the persistent effect of the already acquired statistical representations (Weiss, Schwob, & Lebkuecher, 2019). By presenting two artificial speech streams in smaller alternating blocks, individuals were able to learn both statistical structures underlying the input streams, without using explicit contextual cues (e.g., change in speaker) denoting the transitions across streams (Zinszer & Weiss, 2013). However, if statistically incongruent, interfering statistical structures determined the stimuli, the formation of multiple representations was limited (Weiss et al., 2009). Importantly, if individuals were exposed to the second statistical structure immediately after learning had occurred on the first structure presented for a restricted time, both structures were learned. In addition, the different contextual cues did not further enhance performance (Bulgarelli & Weiss, 2016). Altogether, it seems that overlearning the first statistical structure and low variability in how the statistical structures are presented could decrease the attention paid to the input stream, thereby deteriorating the acquisition of the new structure (Bulgarelli & Weiss, 2016). This explanation might be feasible in the case of our findings; however, it should be noted that these studies tested the transition(s) between different statistical structures in the linguistic domain, while our study investigated the unsigned transition from the presence to the absence of a structure in the visuomotor domain. Furthermore, the 2^nd^ order nonadjacent transitional probabilities applied in the present task differed in structure and complexity from those transitional probabilities applied earlier. Nevertheless, we can contribute to this research field by confirming the presence of the primacy effect in a unique multi-context unsupervised learning environment and, this way, by extending the validity of this effect to unlearnable pseudorandom stimuli.

### 4.6 Transfer of prior knowledge

Considering the underlying processes, the present findings raise the question of whether learning transfer has occurred across the task halves. Studies testing the transfer (generalization) of perceptual and motor knowledge usually compare performance observed during a training task with performance observed during a similar testing condition, such as in a familiar task with new parameters or in a related but novel task. Successful transfer occurs if the experience gathered on the training task appears as a performance gain on the novel task (e.g., Dorfberger, Adi-Japha, & Karni, 2012; Karni, 1996; Karni & Bertini, 1997; Korman, Raz, Flash, & Karni, 2003).

In the statistical-sequence learning literature, the implicit transfer of both the perceptual and the motor sequence was shown in a version of the ASRT task that, in the testing phase, included a novel, previously unpracticed alternating motor or perceptual sequence with the same type of stimuli (Hallgató et al., 2013; Nemeth et al., 2009). In a deterministic SRT task, the 2^nd^ order transitional probability structure was implicitly transferred from the training to the testing phase, where the perceptual features of the stimuli differed (i.e., 1^st^ order structure: locations arranged horizontally or according to a square, Huang et al., 2017). Likewise, experiments using the artificial grammar learning task found the implicit transfer of sequential dependencies to novel vocabularies (Tunney & Altmann, 2001). Relatedly, unconscious within- and between-modalities transfer of artificial grammars was shown between training and test strings that changed at the surface level (letters in different vocabularies, notes, symbols) but remained structurally the same (Scott & Dienes, 2010). Another line of research found learning transfer in both directions between different types of memory tasks (motor skill task and word list task) via the extraction of high-level relations between the elements (Mosha & Robertson, 2016).

In contrast to these studies, the present experiment followed a different design: The surface of the stimuli remained the same across the two task phases, but their overall underlying structure changed. In this sense, the observed effect might not be considered as a classical perceptual-motor transfer effect. Indeed, it more likely captures a cognitive transfer effect: After a short experience with the unstructured sequence, participants completing the structured blocks first might have implicitly identified the features common (i.e., triplets) across the two task halves, which might have supported the generalization of the acquired transitional probability structure (cf., Qian et al., 2012; Robertson, 2018; Winkler & Cowan, 2005). Thus, the triplets as the critical building blocks of the structure might have been implicitly recalled in a later phase of task solving, even if their frequencies had changed. On the contrary, participants completing the unstructured blocks first only perceived the relations of stimuli primarily as triplets when they completed the ASRT sequence with biased probabilities. This would be in line with our recent findings that sensitivity to multiple regularities in the ASRT task seems to be grounded in the implicit extraction of the triplet-level probability structure (e.g., Kóbor et al., 2019; Szegedi-Hallgató, Janacsek, & Nemeth, 2019). Together with the previously described various transfer effects, the implicitly acquired prior knowledge seems to be robust to changes in both the surface and the underlying structure of the stimuli.

Related to the present task, an alternative experimental design with biased triplet probabilities would unequivocally test the transfer of the acquired implicit knowledge. For instance, one might use the ASRT sequence as the “structured sequence” and create another “less structured” sequence by keeping the biased distribution of high-(62.5%) and low-probability (37.5%) triplets but omitting the alternating regularity. In a similar between-subjects design, by presenting either the structured or the less structured sequence in the first half of the task and the other sequence in the second half, a future study might compare triplet learning between these two sequences. To follow previous experiments testing the transfer effect, the stimuli in the second half of the task might be different at the surface level (e.g., arrows or colors instead of horizontally arranged positions), but this is not necessary (cf. Gebhart et al., 2009). In another version of this experiment, the less structured sequence could be presented in both task halves for one of the groups, while the structured sequence followed by the less structured sequence could be presented for the other group. In this version, stimuli should differ at the surface level in the second half of the task.

If triplet learning occurs in both sequences, but only after participants completed the structured sequence first followed by the less structured one, it will indicate that knowledge of the transitional probability structure has been transferred across the task halves. However, for participants completing the less structured sequence first or completing only less structured sequences in both halves, we assume that a modest degree of triplet learning would also occur on these less structured sequences because of the initial sensitivity to the triplet-level probability structure (see above). This needs to be tested in additional experiments.

### 4.7 Methodological considerations

From a methodological point of view, it is not obvious how one investigates what has been learned about the statistical structure underlying a given sequence. The study of Reed and Johnson (1994) suggests that to appropriately test whether the complex statistical structure per se has been learned, instead of a random testing sequence, one should use training and testing sequences that differ only in 2^nd^ order transitional probabilities but are identical in terms of 1^st^ order transitional probabilities and other simpler statistics (e.g., location frequency, transition frequency, reversal frequency, coverage, and transition usage). By controlling for the latter characteristics of both sequences, it can be ensured that the RT disruption across the sequences is due to acquiring the 2^nd^ order transitional probability structure that changed from the first to the second sequence. It has also been shown that participants would less likely search for underlying structures if a sequence, compared with another, was subjectively perceived as more random, but, according to objective measures, was more structured (Wolford, Newman, Miller, & Wig, 2004). Considering these issues, in the present experiment, we deliberately avoided the use of fully random sequences; and, instead, we applied “equal probability” unstructured sequences, which were more controlled than the former ones. In addition, low-probability triplets being the major constituents of the unstructured sequences might have contributed to regarding these sequences as more random (Teigen & Keren, 2020).

### 4.8 Conclusions

The present experiment provides evidence that under implicit and incidental learning conditions, perceptual and cognitive processing continues to be influenced by a previously acquired predictable transitional probability structure even after that structure is removed. This implies that, due to the persistency of the acquired representations, unpredictable transitional probabilities are automatically processed according to these prior representations. However, after significant exposure to the unpredictable structure, the updating of prior representations becomes evident: Importantly, this process seems to require a longer stretch of time than that of the acquisition. Although the acquired representations are relatively persistent if the predictable structure is experienced first, protracted exposure to the unpredictable structure preceding the predictable one does not constrain the subsequent acquisition. Finally, the study also highlights the importance of carefully constructing the underlying structure of training and testing sequences in the investigation of statistical-sequence learning.

## Acknowledgments

This research was supported by the National Brain Research Program (project 2017-1.2.1-NKP-2017-00002, PI: D. N.); the Hungarian Scientific Research Fund (OTKA FK 124412, PI: A. K., OTKA PD 124148, PI: K. J., OTKA K 128016, PI: D. N.); the IDEXLYON Fellowship of the University of Lyon as part of the Programme Investissements d’Avenir (ANR-16-IDEX-0005 to D. N.); and the János Bolyai Research Scholarship of the Hungarian Academy of Sciences (to A. K. and K. J.). The authors thank Balázs Török for providing custom-written scripts that generated the unstructured sequences, Borbála German for helping in data acquisition, Zsófia Zavecz for providing helpful comments on the experimental design and on an earlier version of this paper, and the reviewers for their illuminating comments and suggestions.

## Footnotes

1 ASRT-studies with effect size measures for the prior knowledge effect were not available. Therefore, when determining the sample size per group, we followed the guidelines set by some of the previous behavioral ASRT-studies (Hallgató, Győri-Dani, Pekár, Janacsek, & Nemeth, 2013; Horváth, Török, Pesthy, Nemeth, & Janacsek, 2019; Nemeth, Hallgato, Janacsek, Sandor, & Londe, 2009; Nemeth & Janacsek, 2011; Nemeth, Janacsek, & Fiser, 2013; Szegedi-Hallgató et al., 2017; Vékony et al., 2020). On average, the sample size in these studies was approximately 23 per group (*SD* = 10.6).

