## Supplementary material for "Perceiving structure in unstructured stimuli: Implicitly acquired prior knowledge impacts the processing of unpredictable transitional probabilities": Supplemetary Material (from another version of this preprint: version 3, 2020-05-04)

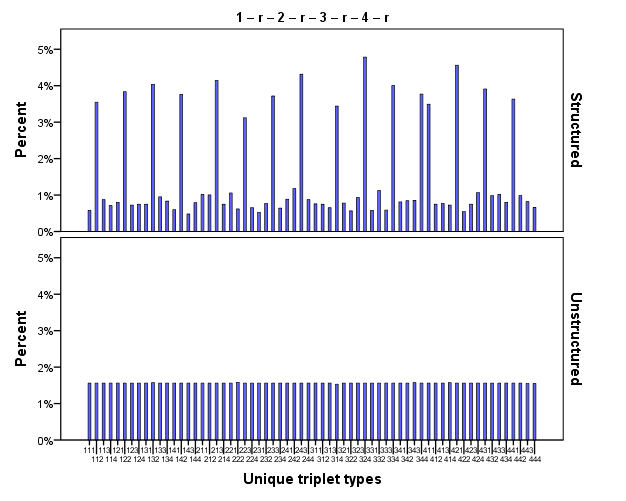

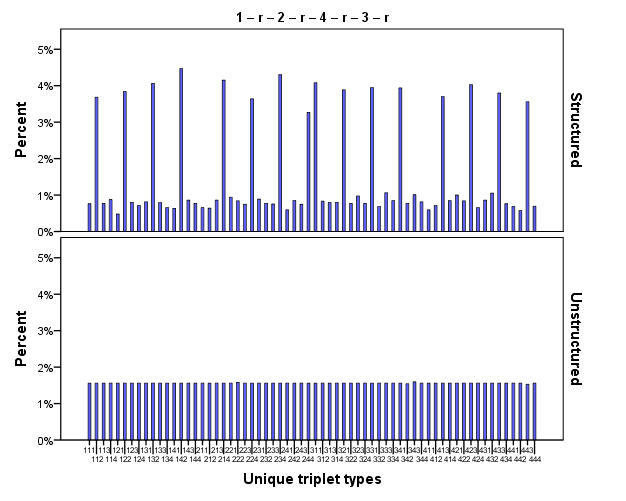

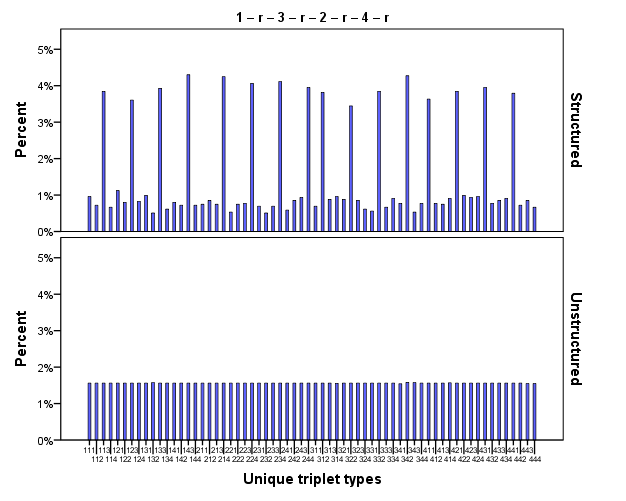

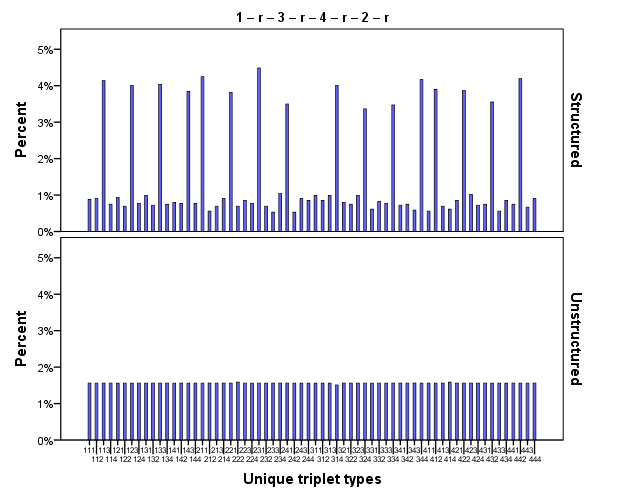

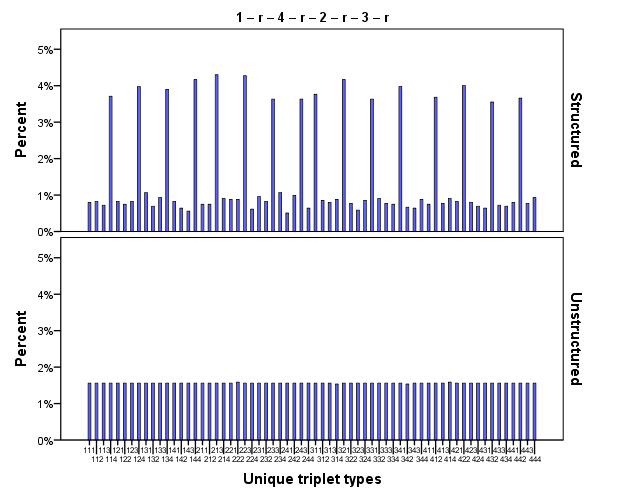

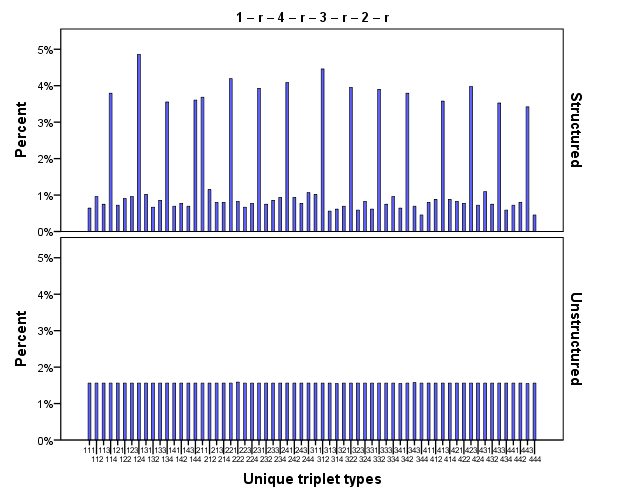

**Figure S1.** Distribution of triplet frequencies for the 64 unique triplet types in the entire sample (*N* = 50) according to the six unique ASRT sequences, separately for the structured and unstructured sequences. In the structured sequences, among the 64 unique triplets, 16 are high-probability triplets and 48 are low-probability ones; the formers are indicated by the higher bins. There is partial overlap across the six unique ASRT sequences in terms of the actual high-probability triplets. The distribution of triplet frequencies is flat in the unstructured sequences, as each triplet occurs with equal probability.

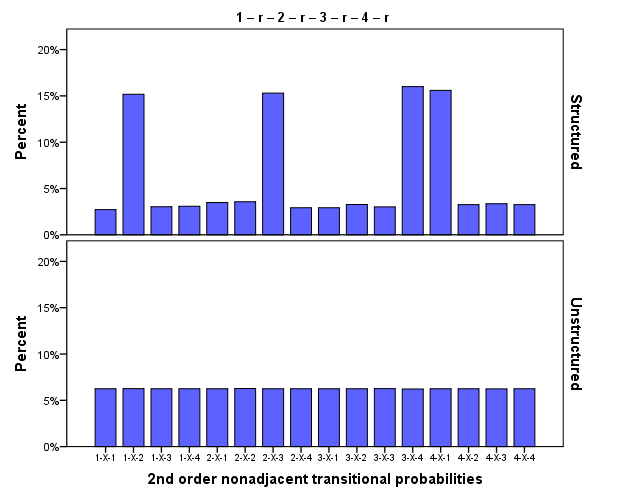

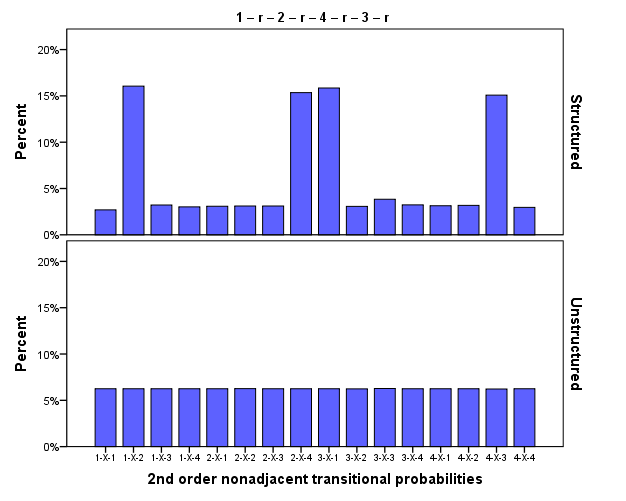

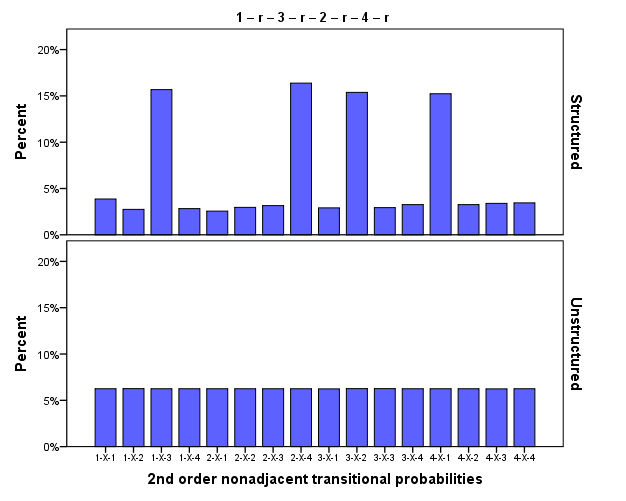

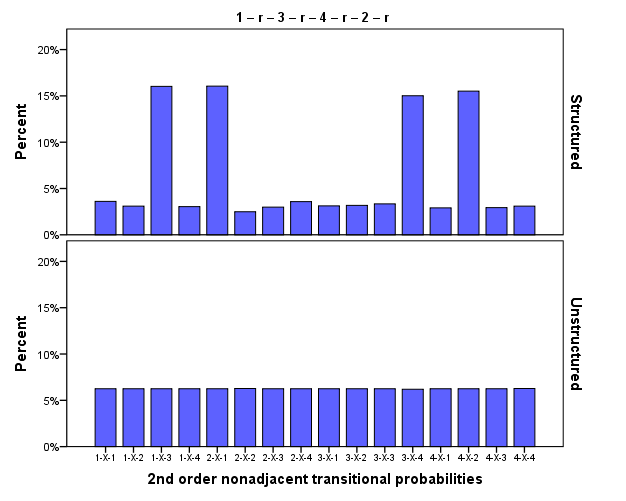

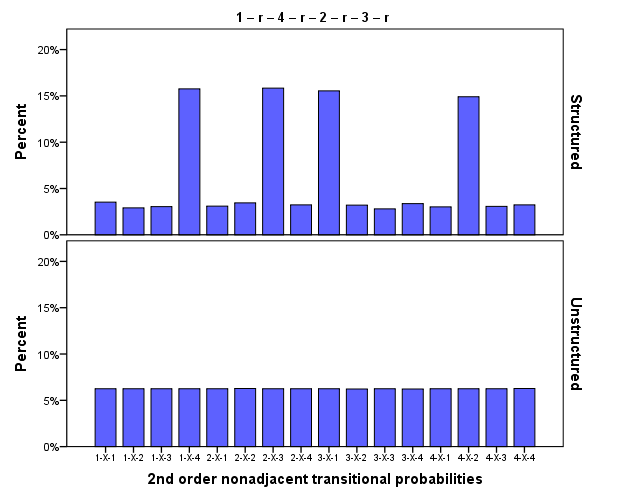

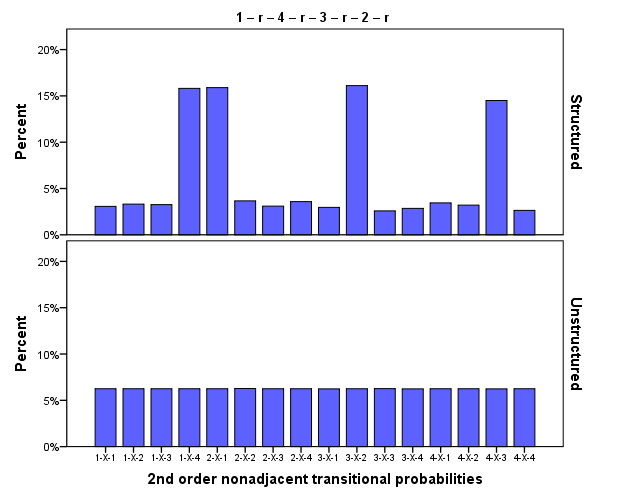

**Figure S2.** Distribution of 2^nd^ order nonadjacent transitional probabilities (16 triplet categories, e.g., 1 – X – 3, 3 – X – 2; X denotes the middle trial of the triplet) in the entire sample (*N* = 50) according to the six unique ASRT sequences, separately for the structured and unstructured sequences. In the structured sequences, four triplet categories include all the high-probability triplets in the given unique ASRT sequence, indicated by the higher bins. There is partial overlap across the six unique ASRT sequences in terms of the high-probability triplet categories. The distribution of 2^nd^ order nonadjacent transitional probabilities is flat in the unstructured sequences, as each of them occurs with equal probability.
